# Pannexin 3 deletion reduces fat accumulation and inflammation in a sex-specific manner

**DOI:** 10.1101/2021.05.04.442670

**Authors:** Charles Brent Wakefield, Vanessa R. Lee, Danielle Johnston, Parastoo Boroumand, Nicolas J. Pillon, Samar Sayedyahossein, Brooke L. O’Donnell, Justin Tang, Rafael E. Sanchez-Pupo, Kevin J. Barr, Robert Gros, Lauren Flynn, Nica M. Borradaile, Amira Klip, Frank Beier, Silvia Penuela

**Affiliations:** Department of Anatomy and Cell Biology, Schulich School of Medicine and Dentistry, University of Western Ontario, London, Ontario, N6A 5C1, Canada; Western’s Bone and Joint Institute, The Dr. Sandy Kirkley Centre for Musculoskeletal Research, University Hospital, London, Ontario, N6G 2V4, Canada; Cell Biology Program, The Hospital for Sick Children, Toronto, Ontario, M5G 0A4, Canada; Department of Biochemistry, University of Toronto, Toronto, Ontario, M5S 1A8, Canada; Department of Physiology, University of Toronto, Toronto, Ontario, M5S 1A8, Canada; Department of Physiology and Pharmacology, University of Western Ontario, London, Ontario, N6A 5C1, Canada; Robarts Research Institute, Schulich School of Medicine and Dentistry, University of Western Ontario, London, Ontario, N6A 5C1 Canada; Department of Chemical and Biomedical Engineering, University of Western Ontario, London, Ontario, N6A 5C1, Canada; Department of Oncology, Division of Experimental Oncology, Schulich School of Medicine and Dentistry, University of Western Ontario, London, Ontario, N6A 5C1, Canada

## Abstract

**Background:** Pannexin 3 (PANX3), is a channel-forming glycoprotein that enables nutrient-induced inflammation *in vitro*, and genetic linkage data suggests it regulates body mass index. Here, we characterized inflammatory and metabolic parameters in global *Panx3* knockout (KO) mice in the context of forced treadmill running (FEX) and high fat diet (HFD).

**Methods:** C57BL/6N (WT) and KO mice were randomized to either a FEX running protocol or no running (SED) from 24 until 30 weeks of age. Body weight was measured biweekly, and body composition was measured at 24 and 30 weeks of age. Male WT and KO mice were fed a HFD from 12 – 28 weeks of age. Metabolic organs were analyzed for a panel of inflammatory markers and PANX3 expression.

**Results:** In females there were no significant differences in body composition between genotypes, which could be due to the lack of PANX3 expression in female white adipose tissue, while male KOs fed a chow diet had lower body weight, and lower fat mass at 24 and 30 weeks of age, which was reduced to the same extent as 6 weeks of FEX in WT mice. Additionally, male KO mice exhibited significantly lower expression of multiple pro-inflammatory genes in white adipose tissue compared to WT mice. While on a HFD body weight differences were insignificant, in KO mice, multiple inflammatory genes were significantly differently expressed in quadriceps muscle and white adipose tissue resulting in a more anti-inflammatory phenotype compared to WT mice. The lower fat mass in male KO mice may be due to significantly fewer adipocytes in their subcutaneous fat compared to WT mice. Mechanistically, adipose stromal cells (ASCs) cultured from KO mice grow significantly slower than WT ASCs.

**Conclusion:** PANX3 is expressed in male adult mouse adipose tissue and may regulate adipocyte numbers, influencing fat accumulation and inflammation.

## Introduction

Obesity is caused by excessive fat accumulation and is a major contributor to many co-morbidities including type II diabetes and cardiovascular disease (1). While exercise training and caloric deficit are effective treatments for obesity, many find these interventions difficult to implement and sustain (2). Genetic factors increase susceptibility to weight gain (3), and understanding which genetic factors underlie obesity will assist clinicians in determining the best pharmacotherapeutic options for a given patient.

Pannexins (PANX1, 2, 3) are channel-forming glycoproteins that allow the passage of ions and metabolites for autocrine and paracrine signaling in a variety of cells (4). Previous reports have shown that PANX1 is expressed in adipocytes and has a functional role in immune cell recruitment (5), adipocyte hypertrophy and fat accumulation (6), glucose metabolism (7), and thermoregulation in brown fat (8). Recent evidence suggests that PANX3 may also play a role in adipogenesis and inflammation (9-11). Using a systems approach involving quantitative trait loci mapping and gene expression network analysis, Halliwill and colleagues found that the *Panx3* gene is linked to body mass index in male mice (9). This group also identified *Panx3* as a component of the *homeodomain-interacting protein kinase 2* (Hipk2) gene network which is involved in adipocyte signaling and differentiation (10). These studies provide indirect evidence that PANX3 may be involved in the molecular mechanism of fat accumulation.

A consequence of excessive fat accumulation is inflammation of adipose tissue (12-15). This inflammation is thought to contribute to many comorbidities (14, 16-19). We previously reported that the saturated-fatty acid palmitate activated cell-intrinsic pro-inflammatory programs in isolated muscle cells and concomitantly increased *Panx3* expression (11). Additionally, we demonstrated that PANX3 channels allowed adenosine triphosphate (ATP) release, attracting monocytes towards the muscle cells (11). This would suggest that PANX3 may be a contributor to nutrient-induced skeletal muscle inflammation by acting as a conduit for ‘find me’ signals to immune cells. Lastly, we observed that HFD induced the expression of *Panx3* in adipose tissue (11), which was the first published finding of *Panx3* expression in mouse adipose tissue. However, its role in diet-induced obesity, fat accumulation, inflammation and metabolism has not been investigated.

Considering the evidence above, we sought out to determine the physiological effects of a global deletion of *Panx3* in mice exposed to forced exercise (FEX) and dietary excess (HFD). In males, *Panx3* Knockout (KO) mice had lower body weight and fat mass, but higher lean mass corrected for body weight, and lower inflammation in adipose and quadriceps tissue compared to WT mice to the same extent as 6 weeks of forced treadmill running. This potentially beneficial loss of natural inflammatory gene expression level was still evident when challenged with caloric excess. However, there were minimal differences between female WT and KO mice, highlighting a sex-specific effect of the *Panx3* deletion. This would suggest that the deletion of *Panx3* attenuates fat accumulation and inflammation in males and could become a useful, sex-specific, genetic target to combat obesity and its associated inflammation.

## Materials and Methods

### Animals and ethics

Experiments performed on animals were approved by the Animal Care Committee of the University Council on Animal Care at the University of Western Ontario, London ON, Canada (UWO # 2019-069), and in accordance with relevant guidelines and regulations. *Panx3* KO mice were generated as described previously (20). *Panx3* KO mice were backcrossed with C57BL/6 N mice from Charles River Canada (Saint-Constant, PQ) until a congenic line was obtained (minimum of 10 backcrossed generations). Mice were weaned at 3 weeks of age and fed either a chow diet (6.2% kcals from fat), Western (45% kcals from fat) or a HFD (60% kcals from fat, Test Diet 58Y1) as described in the respective sections. At termination, mice were sacrificed using carbon dioxide. Immediately after, blood was collected via cardiac puncture, adipose, quadriceps, and liver tissues were collected, immediately snap frozen and stored at -80°C.

### Forced exercise (FEX) protocol

At 24 weeks of age (baseline), mice were randomized to either sedentary (SED) or FEX groups. The FEX groups were forced to run on a treadmill (Columbus Instruments, Ohio) for 6 weeks, 1 hour a day, 5 days a week, at a speed of 11 m/s, and a 10º incline. The mice were encouraged to run using a bottle brush bristle and a shock grid at the end of the treadmill as per the animal ethics protocol. Mice were acclimatized for 10 mins before each session, which consisted of being in the treadmill with no belt movement.

### Body composition

Fat and lean mass composition were measured at baseline and 30 weeks of age using a quantitative magnetic resonance (echo-MRI) mobile unit (Avian Facility of Advanced Research, University of Western Ontario, London, ON, Canada) as described previously (6). Measurements were taken in triplicate to verify the results.

### Blood glucose tolerance and plasma analysis

Mice were fasted 4 hours prior to testing. Fasted blood glucose was measured via a glucometer (OneTouch Ultra). Glucose tolerance testing was conducted by administration of 1 g/kg of glucose by intraperitoneal injection, and blood glucose was monitored at 0, 15, 30, 60, and 120 minutes via tail vein puncture. Glucose area under the curve (AUC) was calculated. At sacrifice, blood was collected, and plasma was isolated. ELISAs (ALPCO, NH, USA) were performed following the manufacturer’s protocol for insulin, cholesterol was assessed by CHOD-PAP kit (Roche Diagnostics, Indianapolis, IN), and triglyceride analysis was conducted by Triglycerol/Glycerol kit (Roche Diagnostics, Indianapolis, IN) following manufacturer’s protocols.

### Metabolic cage analysis

Metabolic analysis was assessed using the Comprehensive Lab Animal Monitoring System (CLAMS) with the Oxymax software (Columbus Instruments, Columbus, OH, USA) at the Robarts Research Institute. Mice were individually caged and acclimated for 24 hours prior to measurement of food consumption, water consumption, energy expenditure, volume of oxygen (VO2) and carbon dioxide (VCO2), respiratory exchange ratio (RER), total activity, total ambulatory activity, and sleep duration, as described previously (6).

### RNA extractions, cDNA synthesis and qPCR

Tissue RNA extraction were performed using total RNA isolation (TRIzol) reagent (Life Technologies) and phenol-chloroform phase separation. Samples were homogenized in TRIzol, mixed with chloroform and centrifuged at 12, 000 x g for 15 minutes at 4°C. The aqueous phase containing the RNA was isolated. Isopropanol was added and samples incubated at room temperature for 30 minutes to precipitate the RNA. The extracted RNA was pelleted by centrifugation at 12, 000 x g for 10 minutes at 4°C. The pellets were washed with 70% ethanol, centrifuged at 7,500 x g for 5 minutes, and then again washed with 100% ethanol and centrifuged lastly at 7,500 x g for 5 minutes. The samples were stored in -80°C.

NanoDrop 2000c spectrophotometer (NanoDrop) was used to quantify the extracted RNA concentration and its purity. SuperScript variable input linear output (VILO) kit for complementary deoxyribonucleic acid (cDNA) synthesis (Life Technologies) was used to synthesize cDNA. cDNA synthesis reaction was performed in 10 μL volume to which up to 125 ng/μL RNA was added and a final concentration of 1X VILO Reaction mix and 1X SuperScript Enzyme mix were loaded. The 96-well plate containing the samples were incubated at 42°C for 60 min, then 85°C for 5 min on a C1000 thermal cycler (Bio-Rad), then stored at -20°C.

A reaction with 20 ng of cDNA was used for each reverse transcriptase quantitative polymerase chain reaction (RT-qPCR) along with 1X TaqMan Fast Advanced Master Mix, and predesigned TaqMan probes (Life Technologies) for the following target genes: Arg1 (Mm00475988_m1), Mrc1 (Mm00485148), IL-10 (Mm00439614), Chi3l3 (Mm00657889), Emr1 (Mm00802529_m1), IL-12a (Mm00434165), CCL2 (Mm00441242), Itgax (Mm00498701_m1), Nos2 (Mm00440502_m1), and Tnf (Mm00443258_m1) on a StepOne Plus Real-Time PCR System (Life Technologies). Samples were held at 95°C for 20 seconds, then cycled from 95°C for 1s to 60°C for 20 s for 40 cycles. Gene expression of target genes were normalized to average of the housekeeping genes *Abt1* (Mm00803824_m1), *Hprt* (Mm03024075_m1), and/or *Eef2* (Mm01171435_gH) using the ΔΔCt method. An inflammatory index score was calculated as the ratio of the sum of pro-inflammatory gene expression over the sum of the anti-inflammatory gene expression and reflects the inflammatory status of the tissue.

For analysis of *Panx3* mRNA expression in visceral fat tissue, RNA was extracted using a combination of Trizol and a Qiagen RNeasy mini kit as was previously described (21). m*Panx3* Forward: TTTCGCCCAGGAGTTCTCATC, Reverse: CCTGCCTGACACTGAAGTTG, m*18S* Forward: GTAACCCGTTGAACCCCATT, Reverse: CCATCCAATCGGTAGTAGCG and m*Hprt*. Normalized mRNA expression levels were analyzed using the ΔΔCT method which was calculated using BioRad CFX Manager Software. Aliquots were taken from the reactions, dyed with ethidium bromide and electrophoresed on a 10% agarose gel.

### Protein analysis

Protein lysates were extracted with lysis buffer containing: 1% Triton X-100, 150 mM NaCl, 10 mM Tris, 1 mM EDTA, 1 mM EGTA, 0.5% NP-40 or a RIPA buffer (50 mM Tris-HCl pH 8.0, 150 mM NaCl, 1% NP-40 (Igepal), 0.5% sodium deoxycholate). Each buffer contained 1 mM sodium fluoride, 1 mM sodium orthovanadate, and half of a tablet of complete-mini EDTA-free protease inhibitor (Roche, Mannheim, Germany). Protein was quantified by bicinchoninic acid (BCA) assay (Thermo Fisher Scientific). Protein lysates (40 μg) were separated by 10% SDS-PAGE and transferred onto a nitrocellulose membrane using an iBlotTM System (Invitrogen, USA). Membranes were blocked with 3% bovine serum albumin (BSA) with 0.05% Tween-20 in 1X phosphate buffer saline (PBS) and incubated with anti-mouse PANX3 antibody (1:1000; PANX3 CT-379) (22), and anti-GAPDH antibody (1:1000; Millipore Cat# MAB374). For detection, IRDye® -800CW and -680RD (Life Technologies, USA) were used as secondary antibodies at 1:10,000 dilutions and imaged using a LI-COR Odyssey infrared imaging system (LI-COR Biosciences, USA). Western blot quantification and analysis was conducted using Image Studio™ Lite (LI-COR Biosciences). Positive controls were generated by ectopic expression of PANX3 constructs in human embryonic kidney 293T (HEK293T) cells as described before (6, 22).

### Histological staining and subcutaneous adipocyte measurements

Dorsal skin samples from adult male WT and KO mice (12-months old) on a chow diet were fixed in 10% neutral buffered formalin and subsequently embedded in paraffin. Sections (5 μm) were deparaffinized in xylene, rehydrated in graded alcohols, and washed in PBS. Parallel tissue sections were stained with hematoxylin/eosin. Images were collected using a Leica DM IRE2 inverted epifluorescence microscope. Measurement of adipocyte cellular size (area) and number of measured adipocytes was performed using the analytical software ImageJ (v.1.50i, National Institute of Health, USA) by a blinded assessor. At least three tissue sections from each mouse were analyzed and individual adipocytes with complete boundaries were selected for quantification and counting.

### 3T3-L1 cell culture and adipogenic induction

Mouse embryonic fibroblast pre-adipocyte (3T3-L1) cells were purchased from ATCC and checked for mycoplasma before use. Cells were grown in Dulbecco’s Modified Eagle’s Medium (DMEM) with 4.5 g/L glucose, 1% Pen-Strep, and 10% calf serum (Thermo Fisher Scientific) and cells below passage 10 were included in the studies. Adipogenic media for days 1-2 contained: DMEM with 4.5 g/L glucose (Thermo Fisher Scientific), 10% calf serum (Thermo Fisher Scientific), 1% Pen-Strep, 100μg/mL of isobutylmethylxanthine (IBMX), 390 ng/mL dexamethosone, and 5μg/mL insulin (Sigma Aldrich). Adipogenic media for days 3-4 contained all of the above components without IBMX or dexamethasone. Following day 4, cells were fed every 2-3 days with DMEM + 10% FBS (Thermo fisher Scientific) until differentiation was complete at day 10.

### Adipose-derived stromal cell isolation

ASCs were isolated as described previously (6). WT and KO male mice were fed on the HFD, with the modification of isolating cells from the inguinal adipose depot and cells were filtered through a 100 μm filter to remove debris prior to cell seeding. Fat from up to three mice was pooled together for each separate isolation. Cells were seeded at high density (80 000 cells/cm^2^) and rinsed 24 hours after isolation with sterile PBS and passaged when confluent (approximately 7 days). ASCs were grown in DMEM: Ham’s F-12 (Sigma Aldrich), supplemented with 10% fetal bovine serum and 1% Pen-Strep and growth medium was changed every 2 days. ASCs used for assays were grown to Passage 2.

### Growth curves and adipogenic differentiation of ASCs

ASCs from WT and *Panx3* KO mice were plated in 12 well plates at a seeding density of 10 000 cells/cm^2^. Cell counts were measured in triplicate every other day up until day 7 using an automated cell counter, Countess II (Thermo Fisher Scientific). Cells were fed every other day with DMEM: Ham’s F12 media, 10% FBS, 1% Pen-Strep, (Sigma Aldrich). Adipogenic induction was conducted with WT ASCs plated in 6 well plates at a seeding density of 30 000 cell/cm^2^. Adipogenic media as previously described (23) with the modifications of substituting 1μg/mL Troglitazone and 0.25 mM IBMX(Sigma Aldrich) for days 1-3. Media was changed every other day for 14 days.

### Statistical analysis

Statistical analyses were performed using GraphPad Prism Version 9.20 (GraphPad, San Diego, CA). Outliers were removed from data sets using the outlier test from GraphPad Prism Version 9.20. Normality tests were used to determine similar variation among the groups for fat mass in males. A power analysis was conducted using the male baseline fat mass mean and standard deviation data to determine an appropriate samples size for the 30-week time point analysis. Body weight progression was analysed using a three-way repeated measures ANOVA with genotype x activity x age as factors. Single time point measures between genotypes were analysed using an unpaired t-test. A two-way ANOVA with genotype x activity as factors was used for 30-week time points, and other two-variable analyses. For blood glucose tolerance curves, a three-way factorial ANOVA was used with genotype x activity x time as factors. Data are presented as mean ± standard error (SEM). N indicates number of animals.

## Results

### Male *Panx3* KO mice weigh less, have less fat mass and more lean mass than WT mice to the same extent as 6 weeks of forced exercise

Male and female WT and KO congenic mice were bred, fed *ad libitum* on a normal rodent chow diet, and randomly allocated to either a SED or FEX protocol from baseline to 30 weeks of age (Fig. 1a). Body weights were tracked bi-weekly, and body composition and blood glucose tolerance were analyzed at baseline and at 30 weeks of age. Postmortem, livers, skeletal muscle, and visceral fat were collected for protein and mRNA analysis.

**Figure 1:**
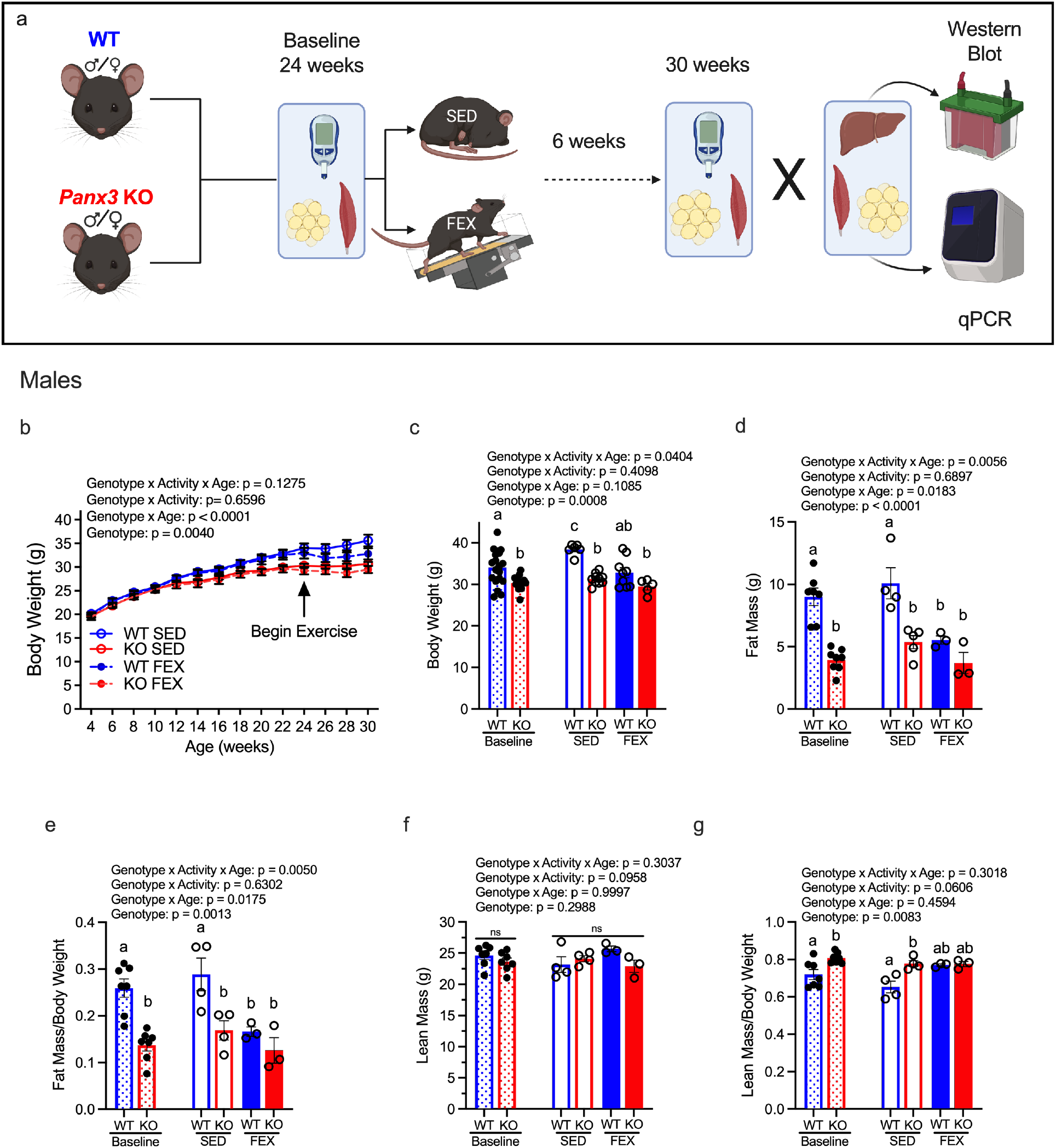
Male *Panx3* KO mice weigh less, have less fat mass and more lean mass than WT mice to the same extent as 6 weeks of forced exercise. Visual graphic of the experimental design. Male and female WT and *Panx3* KO (KO) mice were randomly allocated to either SED or FEX groups from 24 (baseline) to 30 weeks of age (6 weeks) (a). Body weights were measured biweekly, and blood glucose and body composition were measured at baseline and 30 weeks of age. After which, blood and metabolic organs were collected for *ex vivo* analysis. Male body weight development from 4 to 30 weeks of age (b). Baseline comparisons between (checkered bars) KO mice (red, n = 15) and WT mice (blue, n = 19), and at 30 weeks of age for SED (clear bars) and FEX (solid bars) for body weight (c), fat mass (d), fat mass corrected for body weight (e), lean mass (f), and lean mass corrected for body weight (g). Results are expressed as mean ± SEM. A three-way ANOVA was conducted with activity x genotype x age as factors to determine significant differences between the genotypes for each group (n = 3-5). Different letters indicate significantly different groups (p < 0.05). ns: non-significant. WT SED: wildtype sedentary, KO SED: *Panx3* knockout sedentary, WT FEX: wildtype forced exercise, KO FEX: *Panx3* knockout forced exercise.

In males, KO mice weighed significantly less than WT mice as they aged (Fig. 1b). When analyzing the baseline and 30-week body weights with and without exercise, KO mice weighed significantly less at baseline, and at 30 weeks compared to SED WT mice, but were not significantly different from the FEX WTs as exercise attenuated weight gain in WT mice (Fig. 1c).

Considering that *Panx3* may be involved in adipogenesis (10), is linked to body mass index (9), and resulted in lower body weight in the present study, we then sought to determine if this lower body weight in KO mice is due to differences in fat and/or lean mass. KO mice had significantly less fat mass (Fig. 1d) and fat mass corrected for body weight (Fig. 1e) than WT mice at baseline. At 30 weeks of age, SED and FEX KO mice had significantly lower fat mass (Fig. 1d) and fat mass corrected for body weight (Fig. 1e) compared to SED WT mice. Interestingly, FEX significantly decreased fat mass in WT mice, while FEX had no additional effect on fat mass in KO mice (Fig. 1d & e). This suggests that the deletion of *Panx3* alone has a profound effect on fat mass that is not further decreased by FEX. While there were no significant differences in raw lean mass among the groups (Fig. 1f), when lean mass was corrected for body weight, KO mice had significantly more lean mass compared to WT mice at baseline (Fig. 1g). Additionally, at 30 weeks of age, SED KO mice had significantly higher lean mass corrected for body weight compared to SED WT mice (Fig. 1g). However, the FEX WT mice had similar lean mass when corrected for body weight compared to KO mice. These results suggest that the deletion of *Panx3* reduces fat mass and increases lean mass to the same extent as 6 weeks of FEX in male WT mice.

Using individual metabolic cage analysis, we found that there were no significant differences in O2 Volume (Fig. S1a), CO2 Volume (Fig. S1b), Respiratory Exchange Ratio (Fig. S1c), Energy Expenditure (Fig. S1d), Water Consumed (Fig. S1f), Total Activity (Fig. S1g), Ambulatory Activity (Fig. S1h), or Sleep Time (Fig. S1i) between male WT and KO mice. However, there was a main effect for Food Consumption suggesting that KO mice ate more food overall (Fig. S1e), indicating that the reduced body weight and fat mass in KO mice was not due to increased activity or reduced food consumption.

### Male *Panx3* KO mice have lower inflammatory index in quadriceps and visceral fat tissues compared to WT mice

Changes in adiposity and lean mass could be correlated to inflammatory activation of adipose, liver, and skeletal muscle tissue (24). Therefore, we next determined if *Panx3* deletion influences inflammatory gene expression in these metabolic tissues. Liver, quadriceps muscle and visceral white adipose tissues were collected from male WT and KO mice from both SED and FEX groups for analysis of inflammatory genes and an inflammatory index was calculated. Macrophage markers *Emr1* and *Itgax* (CD11c), pro-inflammatory genes *Tnfα, Nos2, Il12a, Ccl2* and *Il6* and anti-inflammatory markers *Arg1, Mrc1, Il10* and *Chi3l3* were analyzed in these tissues using RT-qPCR. An inflammatory index was calculated as the ratio of the sum of the pro-inflammatory markers over the sum of the anti-inflammatory markers and reflects the inflammatory status of the tissue. In quadriceps, SED WT mice had significantly higher *Emr1* compared to all other groups, while there was a main effect for KO mice having significantly higher *Nos2, Tnfα*, and *Il12a* compared to WT mice (Fig. 2a). When analyzing the inflammatory index for the quadriceps, no group was significantly different than SED KO mice (Fig. 2b). In visceral adipose tissue, KO mice had significantly lower pro-inflammatory markers *Emr1, Itgax, Ccl2, Tnfα*, and anti-inflammatory markers *Mrc1* (Fig. 2c). KO mice had significantly lower inflammatory indexes in visceral fat compared to both SED and FEX WT mice (Fig. 2d). However, there were no significant differences among the groups for any genes in the liver or the overall inflammatory index (Fig. 2e & f). These results suggest that the deletion of *Panx3* results in a potential shift of inflammatory tone in skeletal muscle and white adipose tissue, comparable to the effects of FEX.

**Figure 2:**
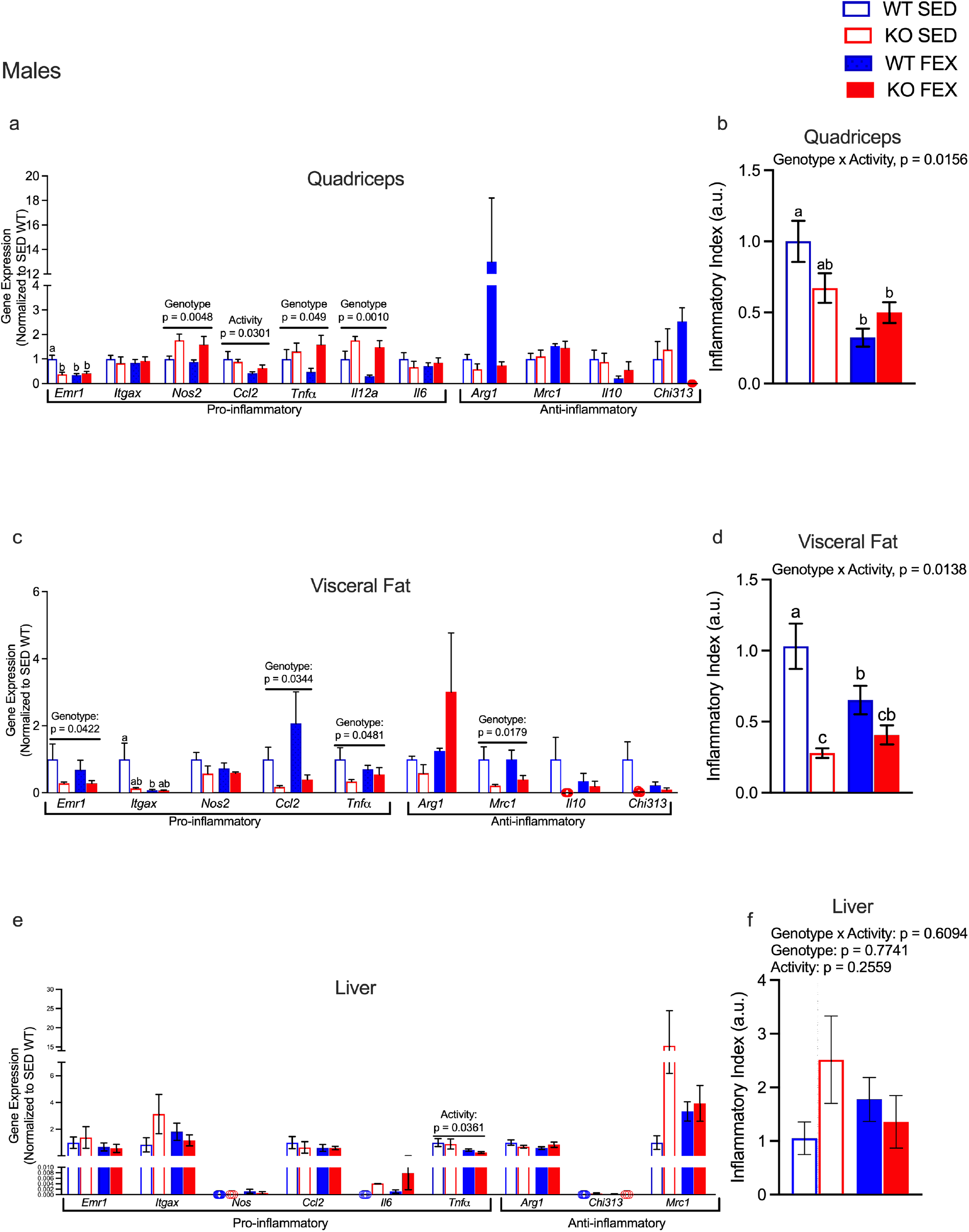
Male *Panx3* KO mice have lower inflammatory index in quadriceps and visceral fat tissues compared to WT mice. WT (blue bars) and KO (red bars) mice were allocated to either SED (clear bars) or FEX (solid bars) groups from 24 (baseline) to 30 weeks of age. Quadriceps (a & b), visceral fat (c & d), and liver (e & f) tissues were collected, and mRNA expression was analyzed by RT-qPCR for macrophage markers *Emr1* and *Itgax* (CD11c), pro-inflammatory genes *Tnfα, Nos2, Il12a, Ccl2* and *Il6* and anti-inflammatory markers *Arg1, Mrc1, Il10* and *Chi3l3*. An inflammatory index score was calculated as the ratio of the sum of the pro-inflammatory over the sum of the anti-inflammatory markers and reflects the inflammatory status of the tissue (b, d, f). A two-way ANOVA was conducted with genotype x activity as factors. n = 3-5. mean ± SEM. Different letters indicate significantly different means (p < 0.05). ns: non-significant. a.u.: arbitrary units. WT SED: wildtype sedentary, KO SED: *Panx3* knockout sedentary, WT FEX: wildtype forced exercise, KO FEX: *Panx3* knockout forced exercise.

Despite these changes in body composition and potential anti-inflammatory effect in skeletal muscle and adipose tissue from a global deletion of *Panx3*, there were no significant differences in blood glucose tolerance between genotypes (Fig. S2a, b, & c). However, when analyzing the blood glucose tolerance curves for the 30-week time point the p–value for genotype approached significance (p = 0.0528) (Fig. S2b). There were no significant differences in other circulating measures of metabolic health such as insulin (Fig. S2d), cholesterol (Fig. S2e), and triglycerides (Fig. S2f).

When analyzing circulating measures of inflammation, we found that SED KO mice had significantly lower levels of total adiponectin compared to SED WT mice (Fig. S2g). However, there were no significant differences in heavy molecular weight (HMW) adiponectin (Fig. S2h) or the ratio of total/HMW adiponectin (Fig. S2i). Additionally, there were no differences in serum amyloid A (SAA) between genotypes (Fig. S2j). However, there was a significant main effect for Genotype suggesting KO mice have significantly lower circulating levels of IL-6 compared to WT mice regardless of activity (Fig. S2k).

### Female *Panx3* KO mice weigh slightly less than WT mice with no significant differences in body fat or lean mass

To determine if these differences between genotypes are seen in females, we next compared WT and KO female mice under both SED and FEX conditions. Interestingly in females, KO mice weighed significantly less as they aged, indicated by a significant Genotype x Age interaction (p < 0.0001) (Fig. 3a). When analyzing baseline and 30-week-old data, there was a significant main effect for Genotype, suggesting female KO mice weighed significantly less than WT females, however the effect size was much smaller than in males (Fig. 3b). Despite differences in body weight, there were no significant differences between genotypes in body fat (Fig. 3c), body fat corrected for body weight (Fig. 3d), lean mass (Fig. 3e), or lean mass corrected for body weight (Fig. 3f) at baseline or with and without FEX at 30 weeks of age between genotypes, but there was an effect for Activity in reducing fat mass regardless of genotype. This would suggest that *Panx3*’s role in fat accumulation in females is not as pronounced as in male mice.

**Figure 3:**
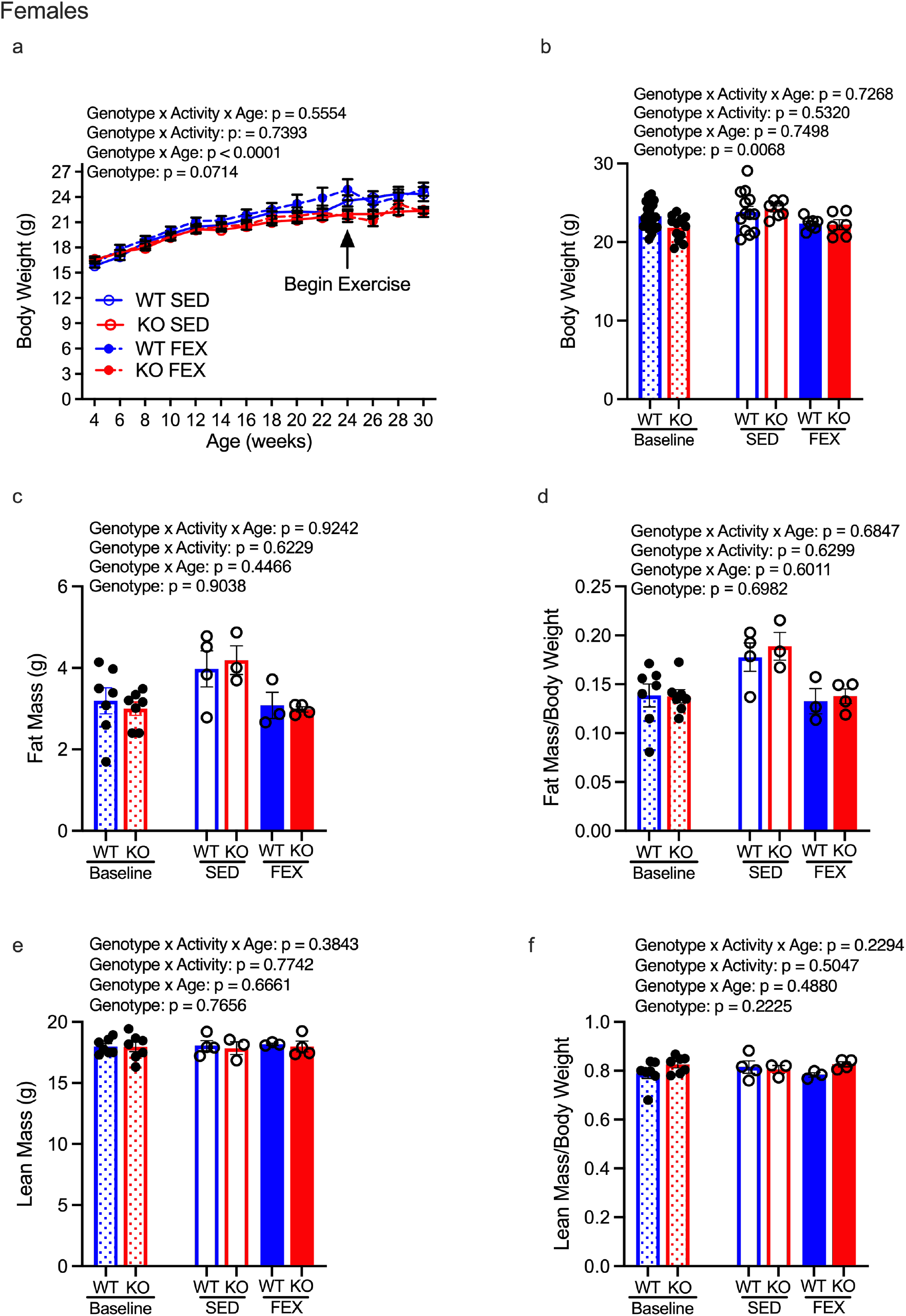
Female *Panx3* KO mice weigh less than WT mice with no significant differences in body fat or lean mass. Female WT (blue) and KO (red) mice were randomly allocated to either SED (clear bars) or FEX (solid bars) group from 24 (baseline) to 30 weeks of age (6 weeks). Body weights were measured biweekly and body composition was measured at baseline and 30 weeks of age. Female body weight measurements from 4–30 weeks age (a). Female body weight comparison at baseline (checkered bars) and 30 weeks of age (N = 7-13) (b). Fat mass (c), fat mass normalized to body weight (d), lean mass (e), and lean mass normalized to body weight (f) was determined by echo-MRI. Results are expressed as mean ± SEM. An unpaired t-test was conducted to assess significant differences between genotypes at baseline of age (N= 7-8). A three-way ANOVA was conducted with activity x genotype x age as factors to determine significant differences between the genotypes (n = 3-4). WT SED: wildtype sedentary, KO SED: *Panx3* knockout sedentary, WT FEX: wildtype forced exercise, KO FEX: *Panx3* knockout forced exercise. ns: non-significant. Different letters indicate significantly different from each other (p < 0.05).

### Female *Panx3* KO mice have higher inflammation in quadriceps and liver tissues compared to WT mice

Female WT and KO mice from both SED and FEX groups were sacrificed, and skeletal muscle, visceral adipose, and liver tissues were excised for analysis as described above. In quadriceps, FEX KOs had significantly higher expression of *Nos2* expression compared to SED WT and KO mice, while WT mice had significantly higher expression of *Mrc1* expression (Fig. 4a). KO mice had significantly higher overall inflammatory index in quadriceps compared to WT mice (Fig. 4b). In visceral fat, both SED and FEX KO mice had significantly lower expression of *TNFα* compared to SED WT mice and significantly higher expression of *Arg1* regardless of activity (Fig. 4c). However, there were no significant differences among the groups when assessing the inflammatory index for visceral fat (Fig. 4d). In liver, KO mice had significantly higher expression of *TNFα* regardless of activity group, and lower *Arg1* and *Chi313* compared to SED WT animals (Fig. 4e). For the overall liver inflammatory index, KO mice had a significantly higher score compared to WT mice regardless of activity (Fig. 4f). This would suggest that in female mice the deletion of *Panx3* leads to a higher inflammatory tone in the quadriceps and liver tissues.

**Figure 4:**
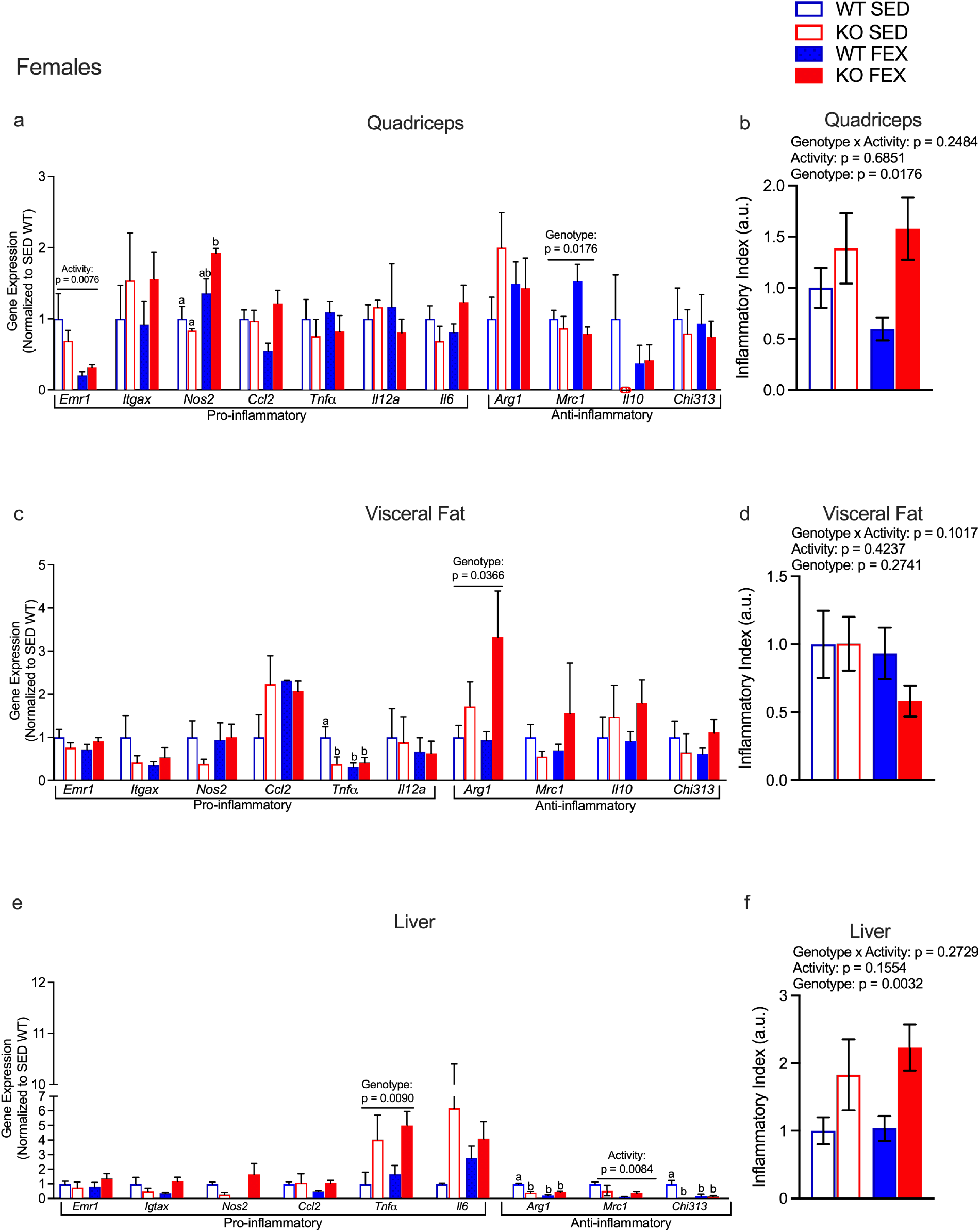
Female *Panx3* KO mice have higher inflammatory index in quadriceps and liver tissues compared to WT mice. WT (blue) and KO (red) mice were allocated to either SED (clear bars) or FEX (solid bars) groups from 24 to 30 weeks of age. Quadriceps (a & b), visceral fat (c & d), and liver (e & f) tissues were collected, and mRNA expression was analyzed by RT-qPCR for macrophage markers *Emr1* and *Itgax* (CD11c), pro-inflammatory genes *Tnfα, Nos2, Il12a, Ccl2* and *Il6* and anti-inflammatory markers *Arg1, Mrc1, Il10* and *Chi3l3*. An inflammatory index score was calculated as the ratio of the sum of pro-inflammatory over the sum of anti-inflammatory markers and reflects the inflammatory status of the tissue (b, d, f). A two-way ANOVA with genotype x activity as factors was conducted. n = 3-5. ns: not significant, mean ± SEM. arbitrary units (a.u.). Different letters indicate significantly different means (p < 0.05). WT SED: wildtype sedentary, KO SED: *Panx3* knockout sedentary, WT FEX: wildtype forced exercise, KO FEX: *Panx3* knockout forced exercise.

When assessing blood glucose tolerance in females, there was no significant difference at baseline (Fig. S3a), however at the 30-week timepoint there was a significant Genotype x Activity interaction (Fig. S3b). While there was no significant effect of exercise on AUC in KO mice, WT mice seemed to have improved glucose handling with FEX (Fig. S3c). Additionally, there was an overall main effect of Genotype for AUC, suggesting KO mice have improved glucose handling (Fig. S3c). There were no significant Genotypic effects on insulin (Fig. S3d), triglycerides (Fig. S3e), or cholesterol (Fig. S3f). However, there was a main effect of Activity for insulin (Fig. S3d).

When assessing circulating levels of inflammatory markers there were no significant differences in total adiponectin (Fig. S3g), HMW adiponectin (Fig. S3h), total/HMW adiponectin (Fig. S3i), SAA (Fig. S3j), and IL-6 (Fig. S3k).

### PANX3 expression is higher in male visceral fat and is regulated by FEX and dietary caloric excess

Considering *Panx3* deletion is producing sex differences in fat and lean mass, we next wanted to determine if PANX3 expression is different between male and female visceral fat. Protein from visceral fat of both SED and FEX male and female WT mice was isolated and ran on a Western blot (Fig. 5a & b). Males had significantly higher expression of PANX3 compared to females regardless of activity levels. Interestingly, FEX seemed to increase PANX3 expression, but this did not reach significance (p = 0.0607) (Fig. 5a). Next, considering Pillon *et al*. previously showed that HFD significantly increases *Panx3* mRNA expression in fat (11), we wanted to confirm these results, and determine if FEX was able to counter this effect. Visceral fat mRNA was isolated from male WT mice that ate either chow or a Western diet (45% kcal from fat) and were subjected to either SED or FEX conditions (Fig. 5c & d). Western diet significantly increased *Panx3* expression compared to chow fed animals, however, FEX attenuated this expression in Western fed animals (Fig. 5c & d). Next, we fed male WT and KO mice a HFD (60% kcal from fat) from 12 to 28 weeks of age. Like Pillon *et al*. (11) and our mRNA results, HFD significantly increased PANX3 expression in fat compared to chow fed animals (Fig. 5e & f). Next, we wanted to determine if deleting *Panx3* would have a protective effect on body weight under HFD feeding as seen in chow fed male mice. However, there were no significant differences in raw body weight (Fig. 5g) or body weight fold change (Fig. 5h) between WT and KO mice when on a HFD.

**Figure 5:**
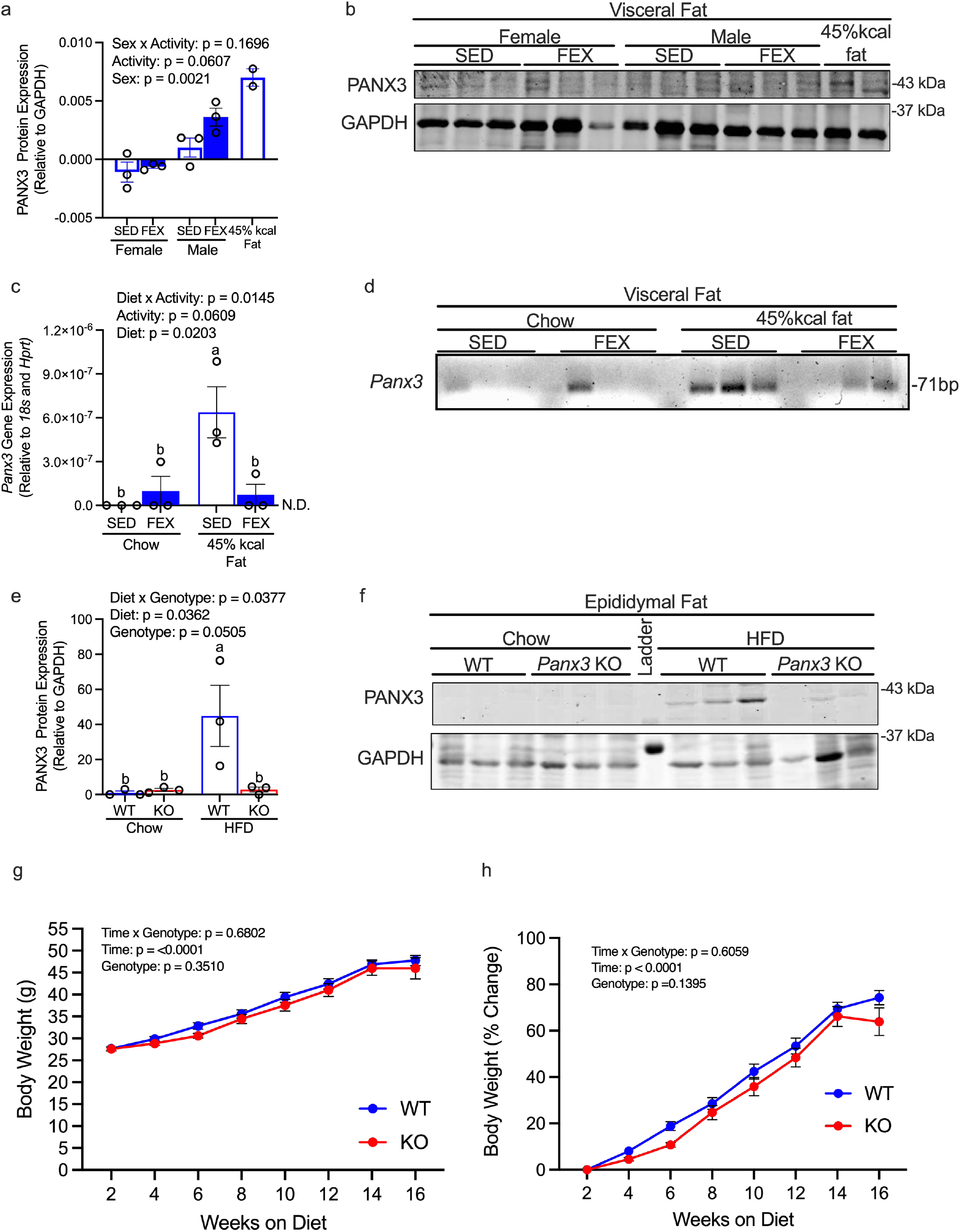
PANX3 expression is higher in male visceral fat tissue compared to females, and is regulated by FEX and dietary fat intake. Male and female WT mice were fed a normal chow diet and allocated to either SED or FEX groups, and their visceral fat was isolated and analysed for PANX3 protein expression (a & b). Protein from animals fed a Western diet (45% kcal from fat) was used as a positive control. mRNA from visceral fat of male mice fed a chow or Western diet and subjected to either the SED or FEX protocol was analysed for *Panx3* expression (c & d). Male WT and *Panx3* KO (KO) mice were fed a high fat diet (HFD, 60% kcal from fat) from 12 to 28 weeks of age and epididymal fat was analysed for PANX3 protein expression (e & f). KO mouse tissues were used as a negative control (e & f). ns: not significant, N = 3, n = 3. Different letters indicate significantly different means (p < 0.05). GAPDH was used as a loading control for Western blots, while *18s* and *Hprt* was used for housekeeping genes for qPCR. Body weight (g) and body weight % change (h) was measured in male mice to determine differences in weight gain between genotypes on a HFD. A two-way repeated measures ANOVA with genotype x age was conducted (N = 13–16). Results are expressed as mean ± SEM.

### Male *Panx3* KO mice fed a HFD have less inflammation in epidydimal adipose and skeletal muscle tissue

Considering HFD regulated PANX3 expression in adipose tissue, and PANX3 may mediate nutrient-induced inflammation (11), we then set out to determine if KO mice are protected from diet-induced inflammation. At sacrifice, HFD fed KO and WT mice had liver, quadriceps muscle and epidydimal white adipose tissues (eWAT) collected for analysis of inflammatory markers, as described above. KO mice had significantly higher expression of anti-inflammatory genes *Arg1* and *Il10* compared to WT mice (Fig. 6a), resulting in a significantly lower inflammatory index in quadriceps (Fig. 6 b). In eWAT tissue (Fig. 6 c & d) KO mice had lower expression of pro-inflammatory genes *Ccl2* and *Il6*, and significantly lower *Arg1* and higher *Chi313* anti-inflammatory expression, resulting in significantly lower inflammatory index (Fig. 6d). However, there were no significant differences in individual gene expression (Fig. 6e) or overall inflammatory index in the liver (Fig. 6f) between genotypes. These results suggest that male KO mice have lower skeletal muscle and fat tissue inflammatory tone compared to WT mice while on an HFD.

**Figure 6:**
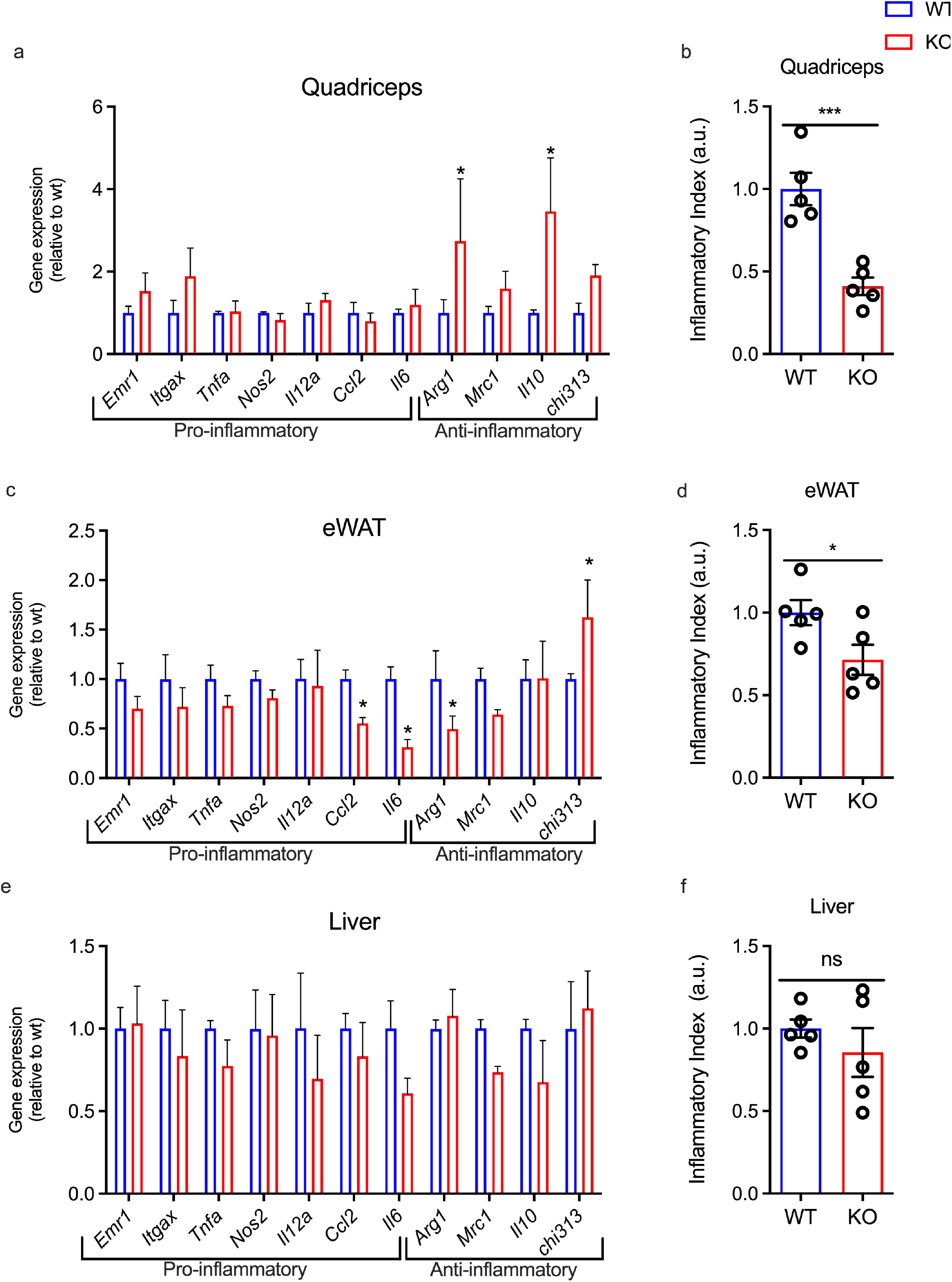
Male *Panx3* KO mice are protected from HFD induced inflammation compared to WT mice. WT and *Panx3* KO (KO) mice were fed a HFD (60% kcal from fat) from 12 to 28 weeks of age. Quadriceps (a & b), epididymal white fat (eWAT) (c & d), and liver (e & f) tissues were collected, and mRNA expression was analyzed by RT-qPCR for macrophage markers *Emr1* and *Itgax* (CD11c), pro-inflammatory genes *Tnfα, Nos2, Il12a, Ccl2* and *Il6* and anti-inflammatory markers *Arg1, Mrc1, Il10* and *Chi3l3*. An inflammatory index score was calculated as the ratio of the sum of pro-inflammatory over the sum of the anti-inflammatory markers and reflects the inflammatory status of the tissue. An unpaired t-test was conducted to determine significant differences between genotypes. N = 5, * = p < 0.05. Results are expressed as mean ± SEM. ns: non-significant, arbitrary units (a.u.).

### *Panx3* KO mice have fewer adipocytes, and their ASCs grow slower than WT mice

Considering we found that male KO mice have significantly lower fat mass than WT mice, we wanted to determine if this was the result of less adipocytes or a reduction in adipocyte hypertrophy. While there were no differences in the size of subcutaneous adipocytes between KO and WT mice (Fig. 7a & b), KO mice had significantly fewer adipocytes (Fig. 7a & c). This suggests that the deletion of *Panx3* may reduce the total number of adipocytes in subcutaneous tissue.

**Figure 7:**
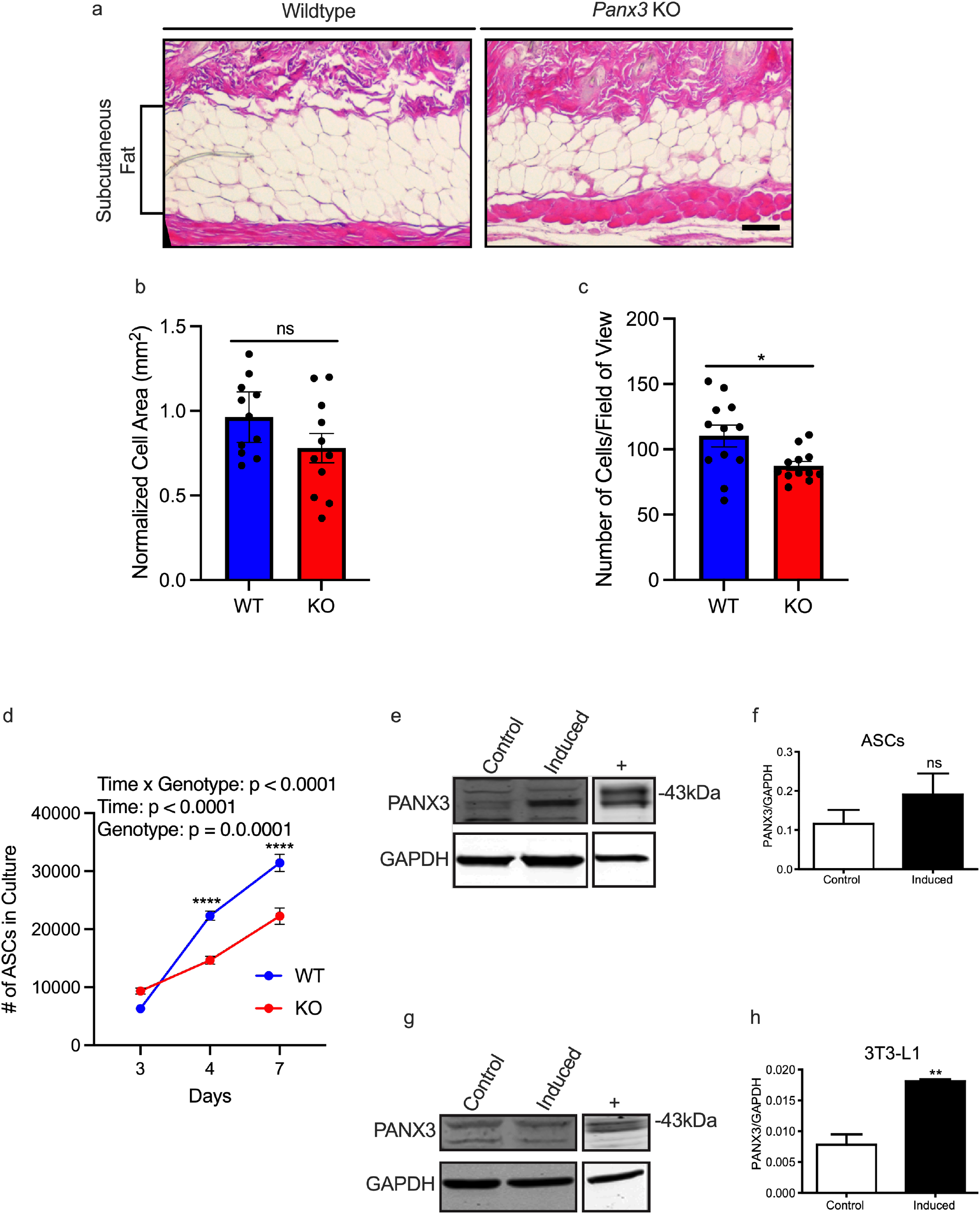
*Panx3* KO mice have fewer adipocytes and their primary adipose stromal cells (ASCs) grow slower than those isolated from WT mice. Representative images of the subcutaneous fat of male WT and *Panx3* KO mice (Scale bar = 100 μm) (a). Adipocyte size (normalized to WT size) (b) and the number of cells (normalized to the standardized area of view) (c) were quantified. ASCs were isolated from WT and *Panx3* KO mice and placed in growth media (d). Western blot and quantification showing PANX3 protein expression in ASCs under controlled and induced conditions (for adipocyte differentiation) (e & f). PANX3 expression in terminal differentiated 3T3-L1 pre-adipocytes as shown by Western blots of 3T3-L1 cells cultured under controlled and induced conditions (g) and the quantification of PANX3 protein expression (h). N = 3, n = 3, p < 0.05. Results are expressed as mean ± SEM. * p < 0.05, ****p < 0.0001. ns: non-significant. Overexpressing HEK293 cells were used as positive controls (+).

Considering there is no published literature on PANX3’s role in adipose-derived stromal cells (ASCs) or early adipocyte development, we wanted to determine what role *Panx3* may be playing in cell proliferation and viability. ASCs were isolated from the inguinal fat pads of WT and KO male mice on a HFD, as described previously (6). ASCs cultured from KO mice grew significantly slower (Fig. 7d) at 4 and 7 days compared to WT ASCs. When ASCs were cultured to induce differentiation to adipocytes, there was a non-significant trend for PANX3 protein expression to increase (Fig. 7e & f). In a pre-adipocytes cell line (3T3-L1), PANX3 significantly increased during induction to terminal adipocyte differentiation (Fig. 7g & h). These results suggest that *Panx3* deletion reduces total fat cell number in adult male mice, reduces ASC growth, and may be involved in adipocyte development as its expression is increased during induction.

## Discussion

A number of studies have shown that *Panx3* has a role in the development and pathophysiology of skin (9, 25-28), bone (25, 29, 30), and cartilage (20, 31, 32), and there have been indirect reports of its involvement in body mass index (9) and adipogenesis (10). In previous publications we examined weight and fat mass differences between WT and *Panx3* KO mice at 12 weeks of age (20) or at later ages (18- and 24- months) (32), and we saw no significant differences between genotypes. In this study we observed large significant differences in weight and fat mass in male KO mice at 24 and 30-weeks of age on a chow diet. We also showed that diet and exercise are regulators of PANX3 expression in mouse adipose tissue, and it is significantly more expressed in male adipose tissue. While there was a genotype effect for female KO mice to weigh less at later time points, the deletion of *Panx3* resulted in much larger weight reductions in males. Most of this body weight difference can be accounted for by the lower fat mass in KO mice. This lower fat mass was to the same extent as 6 weeks of FEX in WT mice, however, there were no significant differences in body weight between genotypes when challenged with a HFD. Upon further investigation to determine why *Panx3* deletion may reduce fat mass, we found that these mice have a reduced number of adipocytes in their subcutaneous fat. Furthermore, KO mice had lower levels of multiple pro-inflammatory genes in white adipose and skeletal muscle tissue under both regular chow and HFD feeding. These results suggest that PANX3 is expressed at higher levels in male adipose tissue, and may regulate adipocyte cell proliferation, body fat accumulation and inflammatory gene expression in male mice.

Differences in obesity rates between males and females is the result of a complex interaction between chromosomal, hormonal, gender and behavioural factors (33). While there were significant differences in body weight between WT and KO females as they aged, deleting *Panx3* in males had a much more profound effect on body mass and composition. Considering male C57BL/6 mice are much more susceptible to weight gain and fat expansion under a variety of dietary conditions (34), this may explain why we observed a greater effect in males. Furthermore, quantitative trait loci data linking *Panx3* to body mass index were specific to male mice (9) which further supports the findings in this study. Additionally, we found that PANX3 expression was significantly higher in male adipose tissue compared to females, supporting the notion that *Panx3* plays a role in male but not female adiposity.

Both estrogen and testosterone play a role in metabolic disease and obesity (35). We did not measure sex hormones in this study, and there are no published reports of estrogen and testosterone levels in *Panx3* KO mice. However, PANX3 is expressed in Leydig cells (36) and therefore may influence testosterone production. Interestingly, the female *Panx3* KO mice in this study had significantly higher inflammatory indices in quadriceps and liver tissues. This dichotomy in inflammatory changes between male and female KOs is perplexing, however PANX3 may influence inflammation differently between sexes due to gonadal white adipose tissue, which contributes to differences in lipid metabolism and inflammation between sexes (37).

We have previously shown, in cultured myotubes, that PANX3 propitiates the cell-intrinsic pro-inflammatory effects of the dietary fatty acid palmitate (11). Blocking of PANX3 channels reduced the capacity of cultured skeletal muscle cells to recruit monocytes. While in the present study we did not quantify immune cells, male *Panx3* KO mice had significantly lower expression of *Emr1*, a macrophage marker, in quadriceps and adipose tissue. Additionally, KO mice had reduced expression of pro-inflammatory relative to anti-inflammatory genes, suggesting that the deletion of *Panx3* may attenuate diet and sedentary behaviour induced adipose tissue inflammation (38). While these results support our previous observations regarding PANX3’s role in inflammation, we are unable to determine which cell type is responsible for the altered inflammatory expression.

Exercise has been shown in both animal and humans to have anti-inflammatory effects systemically and in adipose tissue (39-41). Studies in mouse models show that exercise attenuates visceral white adipose tissue inflammation caused by HFD (42), specifically, the recruitment of M1-like macrophages and CD8+ T cells upon exposure to HFD (43). In this study, while we did not assess markers of CD8+T cells, we found that FEX in WT mice resulted in significantly lower levels of macrophage markers in skeletal muscle and adipose tissue. Interestingly, the deletion of *Panx3* also reduced macrophage markers, and resulted in lowering of multiple pro-inflammatory genes that exercise had no effect on. This would suggest that the deletion of *Panx3* has an even greater impact on inflammatory gene expression than 6 weeks of daily FEX.

Previously we reported that *Panx1* KO mice have more fat mass, less lean mass and weigh more than WT mice (6). This suggests an opposite effect than what was observed in the present study with the deletion of *Panx3*. The potential opposing role of *Panx1* and *Panx3* in adipose tissue is not certain, however this may be due to differing functions of the two pannexin isoforms during early development and their involvement in pre-adipocyte fate. While both ASCs from *Panx1* and *Panx3* KOs have reduced proliferation compared to WT ASCs, *Panx1* KO ASCs have enhanced adipogenic differentiation. We did not perform any further assays to assess differentiation fate of *Panx3* KO ASCs, and future research is necessary to study the function of PANX3 in these cells.

Consistent daily exercise is necessary for health, however much of the literature suggests that exercise alone cannot reduce adiposity in people with obesity, and dietary interventions are necessary (44, 45). In mouse models, the extent to which exercise can influence body weight may be dependent on the age, sex, diet, and the nature of the exercise intervention (voluntary versus forced) (46, 47). We found that FEX attenuated weight gain in WT mice because of reduced fat mass and increased lean mass to body weight ratio. FEX had no additional effect on body weight in *Panx3* KO mice however, as these mice do not gain a significant amount of weight or fat mass between 24 and 30 weeks of age. This suggests the presence of the *Panx3* gene is necessary for the natural weight gain that occurs in adult male WT mice under sedentary conditions. What is striking is the magnitude of difference in body weight (difference between means: 7.117g ± 0.6830) and fat mass (difference between means: 4.727g ± 1.238) between genotypes. This equates to an approximately 46.8% reduction in fat mass, which is like the effect of FEX in this experiment.

While we saw drastic effects on body and fat mass from the deletion of *Panx3* in males under SED and regular chow fed conditions, there were no significant differences in body weight during HFD feeding. This finding is in line with multiple previous reports that obesity is mainly the result of excess caloric intake (48, 49). However, we know individuals can vary in how much weight they gain while in a similar caloric excess (50) which would suggest genetic and behavioural factors are also at play. Our findings highlight the importance of taking into consideration environmental and behavioural factors that can interact with genetics when investigating multifactorial diseases such as obesity. Manipulating *Panx3* may not be effective when consuming an excessive caloric diet, however it may be an effective target for patients who are also engaging in healthy caloric consumption.

While *Panx3* levels were low in adipose tissue of chow fed WT animals, it was significantly elevated in mice fed a Western or HFD. This suggests that *Panx3* expression is sensitive to dietary factors, as gleaned from previous work in cell culture models (11). Moreover, FEX was able to counter this diet induced *Panx3* upregulation. This would suggest that exercise is able to inhibit the signalling responsible for PANX3 expression caused by dietary factors. Future studies will be needed to determine what signaling pathways are responsible for the induction and suppression of PANX3 expression by diet and exercise. However, our previous data along with those reported by others indicate that the toll-like receptor 4 (TLR4)/nuclear factor -κB (NF-κB) pathway is activated by the saturated fatty acid palmitate (51). We previously showed that this pathway mediated the expression of *Panx3* mRNA (11). Conversely, moderate aerobic exercise is known to downregulate TLR4, and consequently the proinflammatory NF-κB pathway, thus, potentially inhibiting *Panx3* expression (52).

## Conclusion

We have shown that the deletion of *Panx3* attenuates body weight gain because of lower fat mass in male mice. Additionally, skeletal muscle and adipose tissue of KO mice shift to a more anti-inflammatory phenotype in males. This effect was equivalent to the reduction in body weight gain and fat mass reduction caused by 6 weeks of daily FEX. This suggests PANX3 plays a significant role in fat accumulation and inflammation in adult male mice. This phenotype may be the result of PANX3’s role in adipocyte proliferation in early life. Considering this study used a global KO model, future research is needed to determine if PANX3 functions in other cell types involved in this phenotype. Manipulating PANX3 channel function or expression may be a potential therapeutic target in conjunction with dietary and exercise interventions to manage obesity and associated inflammation in males.

## Author Contributions

CBW project design, mouse husbandry, research data, data analysis, wrote manuscript; VRL research data, data analysis, edited manuscript; DJ research data, reviewed manuscript; PB research data, data analysis; NJP research data, data analysis, edited manuscript; SS, BOD, JT, RESP research data, data analysis, edited manuscript; KJB research data, mouse husbandry, reviewed and edited manuscript; RG research data, metabolic cage analysis; LF research data, edited manuscript; NB research data, edited manuscript; AK research data analysis, manuscript review and editing; FB project design, manuscript review and editing, funding; SP project design and supervision, data analysis, funding, manuscript review and editing.

## Acknowledgements and conflict of interest statement

The authors declare no conflict of interest of any kind with the current manuscript. We thank the funding agencies that supported this work including: Petro-Canada Young Innovator Award – Western University to SP, Ontario Graduate Scholarship to CBW. F.B holds the Canada Research Chair in Musculoskeletal Research and is the recipient of a Foundation Grant from the Canadian Institutes of Health Research (CIHR, Grant #332438). CIHR Foundation Grant (FRN:FDN-143203) to AK. Dr. Silvia Penuela is the guarantor of this work and, as such, had full access to all the data in the study and takes responsibility for the integrity of the data and the accuracy of the data analysis.

## Data availability

All data generated or analyzed during this study are included in the published article (and its online supplementary files). Raw data is available from the corresponding author upon reasonable request.

## References

1. Romieu I, Dossus L, Barquera S, Blottière HM, Franks PW, Gunter M, et al. Energy balance and obesity: what are the main drivers? Cancer Causes Control. 2017;28(3):247–58.

2. Hall KD, Kahan S. Maintenance of lost weight and long-term management of obesity. Med Clin North Am. 2018;102(1):183–97.

3. Thaker VV. Genetic and epigenetics causes of obesity. Adolesc Med State Art Rev. 2017;28(2):379–405.

4. Penuela S, Gehi R, Laird DW. The biochemistry and function of pannexin channels. Biochim Biophys Acta. 2013;1828(1):15–22.

5. Tozzi M, Hansen JB, Novak I. Pannexin-1 mediated ATP release in adipocytes is sensitive to glucose and insulin and modulates lipolysis and macrophage migration. Acta Physiol (Oxf). 2020;228(2):e13360.

6. Lee VR, Barr KJ, Kelly JJ, Johnston D, Brown CFC, Robb KP, et al. Pannexin 1 regulates adipose stromal cell differentiation and fat accumulation. Sci Rep. 2018;8(1):16166.

7. Adamson SE, Meher AK, Chiu YH, Sandilos JK, Oberholtzer NP, Walker NN, et al. Pannexin 1 is required for full activation of insulin-stimulated glucose uptake in adipocytes. Mol Metab. 2015;4(9):610–8.

8. Senthivinayagam S, Serbulea V, Upchurch CM, Polanowska-Grabowska R, Mendu SK, Sahu S, et al. Adaptive thermogenesis in brown adipose tissue involves activation of pannexin-1 channels. Mol Metab. 2020;44:101130.

9. Halliwill KD, Quigley DA, Kang HC, Del Rosario R, Ginzinger D, Balmain A. Panx3 links body mass index and tumorigenesis in a genetically heterogeneous mouse model of carcinogen-induced cancer. Genome Med. 2016;8(1):83.

10. Sjölund J, Pelorosso FG, Quigley DA, DelRosario R, Balmain A. Identification of Hipk2 as an essential regulator of white fat development. Proc Natl Acad Sci U S A. 2014;111(20):7373–8.

11. Pillon NJ, Li YE, Fink LN, Brozinick JT, Nikolayev A, Kuo MS, et al. Nucleotides released from palmitate-challenged muscle cells through pannexin-3 attract monocytes. Diabetes. 2014;63(11):3815–26.

12. Goossens GH. The metabolic phenotype in obesity: fat mass, body fat distribution, and adipose tissue function. Obes Facts. 2017;10(3):207–15.

13. Reilly SM, Saltiel AR. Adapting to obesity with adipose tissue inflammation. Nat Rev Endocrinol. 2017;13(11):633–43.

14. Stolarczyk E. Adipose tissue inflammation in obesity: a metabolic or immune response? Curr Opin Pharmacol. 2017;37:35–40.

15. Unamuno X, Gómez-Ambrosi J, Rodríguez A, Becerril S, Frühbeck G, Catalán V. Adipokine dysregulation and adipose tissue inflammation in human obesity. Eur J Clin Invest. 2018;48(9):e12997.

16. Burhans MS, Hagman DK, Kuzma JN, Schmidt KA, Kratz M. Contribution of adipose tissue inflammation to the development of type 2 diabetes mellitus. Compr Physiol. 2018;9(1):1–58.

17. Hotamisligil GS. Inflammation and metabolic disorders. Nature. 2006;444(7121):860–7.

18. Kusminski CM, Bickel PE, Scherer PE. Targeting adipose tissue in the treatment of obesity-associated diabetes. Nat Rev Drug Discov. 2016;15(9):639–60.

19. Saxton SN, Clark BJ, Withers SB, Eringa EC, Heagerty AM. Mechanistic links between obesity, diabetes, and blood pressure: role of perivascular adipose tissue. Physiol Rev. 2019;99(4):1701–63.

20. Moon PM, Penuela S, Barr K, Khan S, Pin CL, Welch I, et al. Deletion of Panx3 prevents the development of surgically induced osteoarthritis. J Mol Med (Berl). 2015;93(8):845–56.

21. Abitbol JM, Kelly JJ, Barr K, Schormans AL, Laird DW, Allman BL. Differential effects of pannexins on noise-induced hearing loss. Biochem J. 2016;473(24):4665–80.

22. Penuela S, Bhalla R, Gong XQ, Cowan KN, Celetti SJ, Cowan BJ, et al. Pannexin 1 and pannexin 3 are glycoproteins that exhibit many distinct characteristics from the connexin family of gap junction proteins. J Cell Sci. 2007;120(Pt 21):3772–83.

23. Yu G, Wu X, Kilroy G, Halvorsen YD, Gimble JM, Floyd ZE. Isolation of murine adipose-derived stem cells. Methods Mol Biol. 2011;702:29–36.

24. Gao M, Ma Y, Liu D. High-fat diet-induced adiposity, adipose inflammation, hepatic steatosis and hyperinsulinemia in outbred CD-1 mice. PLoS One. 2015;10(3):e0119784.

25. Abitbol JM, O’Donnell BL, Wakefield CB, Jewlal E, Kelly JJ, Barr K, et al. Double deletion of Panx1 and Panx3 affects skin and bone but not hearing. J Mol Med (Berl). 2019;97(5):723–36.

26. Celetti SJ, Cowan KN, Penuela S, Shao Q, Churko J, Laird DW. Implications of pannexin 1 and pannexin 3 for keratinocyte differentiation. J Cell Sci. 2010;123(Pt 8):1363–72.

27. Zhang P, Ishikawa M, Doyle A, Nakamura T, He B, Yamada Y. Pannexin 3 regulates skin development via Epiprofin. Sci Rep. 2021;11(1):1779.

28. Zhang P, Ishikawa M, Rhodes C, Doyle A, Ikeuchi T, Nakamura K, et al. Pannexin-3 deficiency delays skin wound healing in mice due to defects in channel functionality. J Invest Dermatol. 2019;139(4):909–18.

29. Bond SR, Lau A, Penuela S, Sampaio AV, Underhill TM, Laird DW, et al. Pannexin 3 is a novel target for Runx2, expressed by osteoblasts and mature growth plate chondrocytes. J Bone Miner Res. 2011;26(12):2911–22.

30. Ishikawa M, Williams G, Forcinito P, Ishikawa M, Petrie RJ, Saito K, et al. Pannexin 3 ER Ca(2+) channel gating is regulated by phosphorylation at the Serine 68 residue in osteoblast differentiation. Sci Rep. 2019;9(1):18759.

31. Iwamoto T, Nakamura T, Doyle A, Ishikawa M, de Vega S, Fukumoto S, et al. Pannexin 3 regulates intracellular ATP/cAMP levels and promotes chondrocyte differentiation. J Biol Chem. 2010;285(24):18948–58.

32. Moon PM, Shao ZY, Wambiekele G, Appleton C, Laird DW, Penuela S, et al. Global deletion of pannexin 3 accelerates development of aging-induced osteoarthritis in mice. Arthritis Rheumatol. 2021.

33. Link JC, Reue K. Genetic basis for sex differences in obesity and lipid metabolism. Annu Rev Nutr. 2017;37:225–45.

34. Ingvorsen C, Karp NA, Lelliott CJ. The role of sex and body weight on the metabolic effects of high-fat diet in C57BL/6N mice. Nutr Diabetes. 2017;7(4):e261.

35. Shin JH, Hur JY, Seo HS, Jeong YA, Lee JK, Oh MJ, et al. The ratio of estrogen receptor alpha to estrogen receptor beta in adipose tissue is associated with leptin production and obesity. Steroids. 2007;72(6-7):592–9.

36. Turmel P, Dufresne J, Hermo L, Smith CE, Penuela S, Laird DW, et al. Characterization of pannexin1 and pannexin3 and their regulation by androgens in the male reproductive tract of the adult rat. Mol Reprod Dev. 2011;78(2):124–38.

37. Varghese M, Griffin C, McKernan K, Eter L, Lanzetta N, Agarwal D, et al. Sex differences in inflammatory responses to adipose tissue lipolysis in diet-induced obesity. Endocrinology. 2019;160(2):293–312.

38. Winn NC, Cottam MA, Wasserman DH, Hasty AH. Exercise and adipose tissue immunity: outrunning inflammation. Obesity. 2021;29(5):790–801.

39. Paolucci EM, Loukov D, Bowdish DME, Heisz JJ. Exercise reduces depression and inflammation but intensity matters. Biol Psychol. 2018;133:79–84.

40. Petersen AM, Pedersen BK. The anti-inflammatory effect of exercise. J Appl Physiol (1985). 2005;98(4):1154–62.

41. Suzuki K. Chronic inflammation as an immunological abnormality and effectiveness of Exercise. Biomolecules. 2019;9(6).

42. Rosa-Neto JC, Silveira LS. Endurance exercise mitigates immunometabolic adipose tissue disturbances in cancer and obesity. Int J Mol Sci. 2020;21(24).

43. Kawanishi N, Yano H, Yokogawa Y, Suzuki K. Exercise training inhibits inflammation in adipose tissue via both suppression of macrophage infiltration and acceleration of phenotypic switching from M1 to M2 macrophages in high-fat-diet-induced obese mice. Exerc Immunol Rev. 2010;16:105–18.

44. Swift DL, Johannsen NM, Lavie CJ, Earnest CP, Church TS. The role of exercise and physical activity in weight loss and maintenance. Prog Cardiovasc Dis. 2014;56(4):441–7.

45. Swift DL, McGee JE, Earnest CP, Carlisle E, Nygard M, Johannsen NM. The effects of exercise and physical activity on weight loss and maintenance. Prog Cardiovasc Dis. 2018;61(2):206–13.

46. Carpenter KC, Strohacker K, Breslin WL, Lowder TW, Agha NH, McFarlin BK. Effects of exercise on weight loss and monocytes in obese mice. Comp Med. 2012;62(1):21–6.

47. McCabe LR, Irwin R, Tekalur A, Evans C, Schepper JD, Parameswaran N, et al. Exercise prevents high fat diet-induced bone loss, marrow adiposity and dysbiosis in male mice. Bone. 2019;118:20–31.

48. Golay A, Bobbioni E. The role of dietary fat in obesity. Int J Obes Relat Metab Disord. 1997;21 Suppl 3:S2–11.

49. Wang L, Wang H, Zhang B, Popkin BM, D. S. Elevated fat intake increases body weight and the risk of overweight and obesity among chinese adults: 1991-2015 Trends. Nutrients. 2020;12(11).

50. Levine JA, Eberhardt NL, Jensen MD. Role of nonexercise activity thermogenesis in resistance to fat gain in humans. Science. 1999;283(5399):212–4.

51. Ajuwon KM, Spurlock ME. Palmitate activates the NF-kappaB transcription factor and induces IL-6 and TNFalpha expression in 3T3-L1 adipocytes. J Nutr. 2005;135(8):1841–6.

52. Cavalcante PAM, Gregnani MF, Henrique JS, Ornellas FH, Araújo RC. Aerobic but not resistance exercise can induce inflammatory pathways via toll-like 2 and 4: a systematic review. Sports Med Open. 2017;3(1):42.

