## Supplemental Figures and Legends for "Pannexin 3 deletion reduces fat accumulation and inflammation in a sex-specific manner"

#### Supplemental Figure Legends

**Supplemental Figure 1: *Panx3* KO mice eat more food in a metabolic cage but have similar metabolic activity and sleep.**

WT and *Panx3* KO male mice fed *ad libitum* on a normal chow diet were placed in metabolic cages to assess metabolism and activity during their sleeping period (light) and the active period (dark). O2 (a) and CO2 consumption (b), respiratory exchange ratio (c), energy expenditure (d), food consumption (e), water consumption (f), total activity (g), ambulatory activity (h), and sleep time (i). a two-way ANOVA was conducted with genotype x time as factors. n=4. Results are expressed as mean ± SEM.

**Supplemental Figure 2: No overt differences in markers of metabolism or circulating markers of inflammation in male SED or FEX WT and KO mice.**

Male WT and *Panx3* KO (KO) mice were fed a chow diet and allocated to either SED or FEX conditions. Blood glucose curves for 24 weeks (baseline) (a) and 30 weeks for SED and FEX animals (b). N = 3–4. Blood glucose area under the curve (AUC) from baseline and 30 weeks of age (c). A three-way ANOVA with genotype x activity x age as factors was conducted. Plasma insulin (d), cholesterol (e) triglycerides (f), total adiponectin (g), heavy molecular weight (HMW) adiponectin (h), ratio of HMW/total adiponectin (i), serum amyloid A (SAA) (j), and interleukin 6 (*Il6*) (k). A two-way ANOVA with genotype x activity as factors. n = 4–5.

**Supplemental Figure 3: *Panx3* KO female mice have lower blood glucose under SED conditions with no differences in circulating measures of inflammation.**

Female WT and *Panx3* KO (KO) mice were fed a chow diet and allocated to either SED or FEX conditions. Blood glucose curves for 24-week (baseline) (a) and 30-week timepoint for SED and FEX animals (b). Blood glucose area under the curve (AUC) from baseline and 30 weeks of age (c). A three-way ANOVA with genotype x activity x age as factors was conducted. n = 3–4. Plasma insulin (d), triglycerides (e) cholesterol (f), total adiponectin (g), heavy molecular weight (HMW) adiponectin (h), ratio of HMW/total adiponectin (i), serum amyloid A (SAA) (j), and interleukin 6 (IL6) (k). A two-way ANOVA with genotype x activity as factors was conducted. n = 4–5.

Supplemental 1

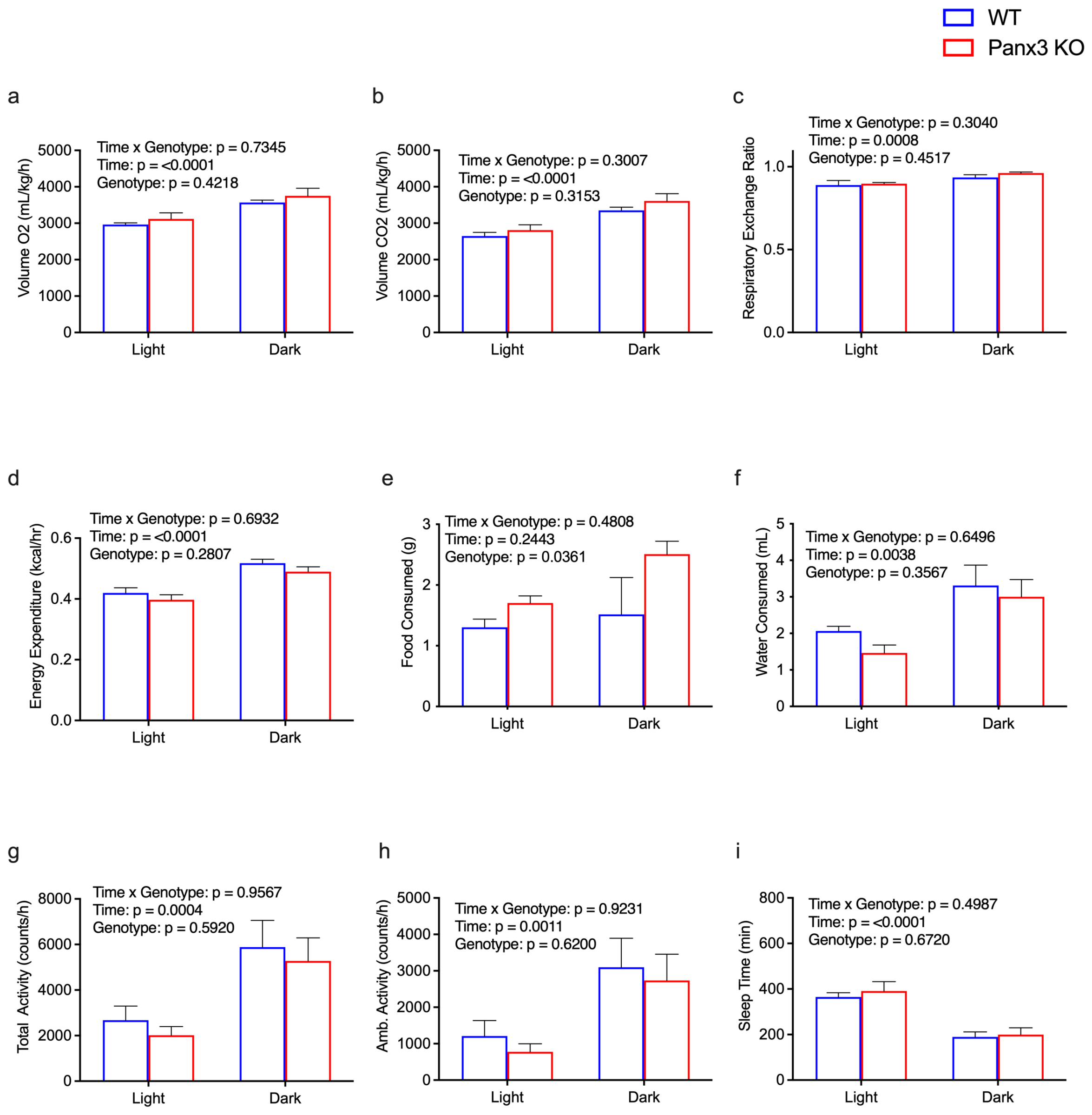

#### Supplemental 2

##### Males

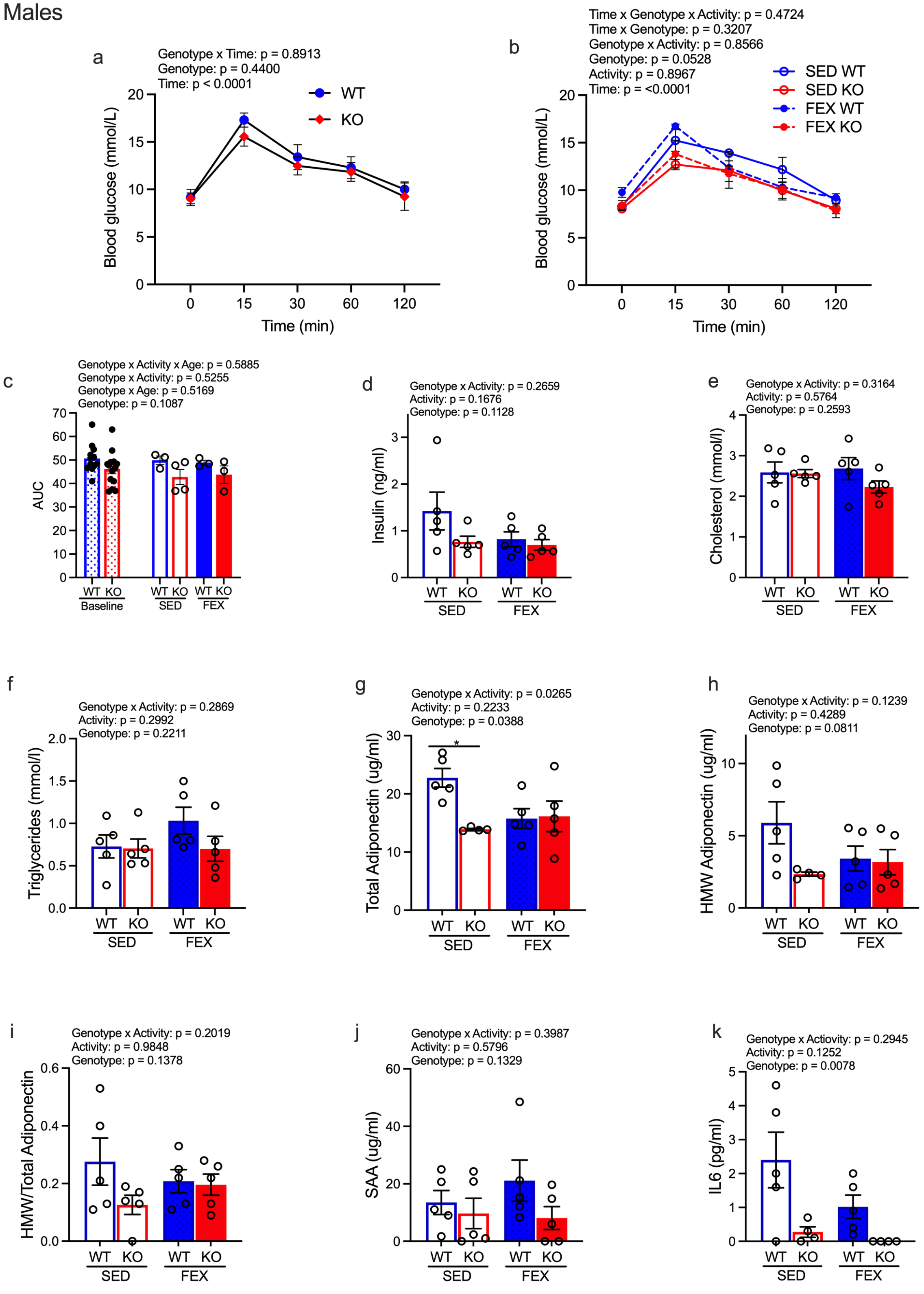

### Supplemental 3

#### Females

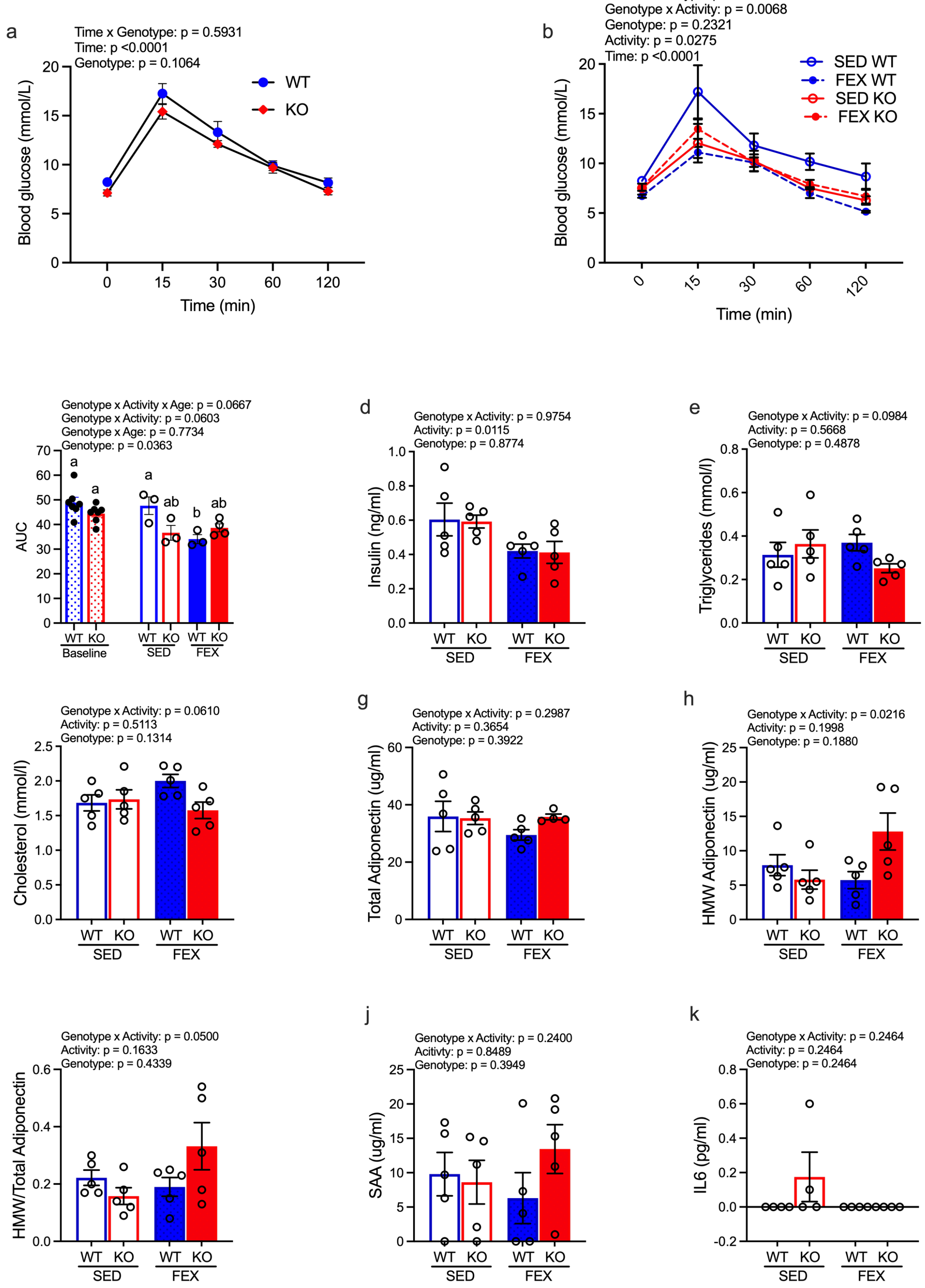
